# Chromatin mobility after DNA damage is modified to enhance long distance explorations and minimize local resampling

**DOI:** 10.1101/042051

**Authors:** Judith Miné-Hattab, Vincent Recamier, Ignacio Izeddin, Rodney Rothstein, Xavier Darzacq

## Abstract

The dynamic organization of genes inside the nucleus is an important determinant for their function. Using ultra-fast microscopy in *S. cerevisiae* cells and improved analysis of mean square displacements, we quantified DNA motion at time scales ranging from 10 milliseconds to minutes and found that following DNA damage, DNA exhibits distinct sub-diffusive regimes. In response to double-strand breaks, chromatin is more mobile at large time scales but, surprisingly, its mobility is dramatically reduced at short time scales. This effect is even more pronounced at the break. Such pattern of dynamics is consistent with a global increase in chromatin persistence length following DNA damage. Scale-dependent nuclear exploration is regulated by the Rad51 repair protein, both at the break and throughout the genome. We propose a model in which stiffening of the damaged ends by the repair complex, combined with global increased stiffness, act like a “needle in a decompacted ball of yarn”, enhancing the ability of the break to traverse the chromatin meshwork.

## INTRODUCTION

The dynamic organization of the nuclear genome is essential for many biological processes and is often altered in cells from diseased tissue (Misteli, 2010). Recent advances in live cell imaging make it possible to visualize the dynamic organization of chromosomal loci inside living nuclei (Bronshtein Berger et al., 2013). In the presence of double-strand breaks (DSB) in *S. cerevisiae* and some mammalian cell lines, DNA mobility is dramatically increased (Chiolo et al., 2011; Dimitrova et al., 2008; Dion et al., 2012; Haber and Leung, 1996; Jakob et al., 2011; Lawrimore et al., 2017; Miné-Hattab and Rothstein, 2012; Neumann et al., 2012; Roukos et al., 2013). In diploid yeast, increased mobility following DSBs likely favors pairing between homologues during repair by homologous recombination (HR) (Miné-Hattab and Rothstein, 2012). In addition, in response to random DSBs in diploid yeast, undamaged loci also exhibit increased mobility albeit to a smaller extent than the damaged locus. Increased mobility of undamaged loci is called global or genome-wide increased mobility (Miné-Hattab and Rothstein, 2012, 2013). In haploid yeast, increased mobility in response to DSBs is thought to promote ectopic repair events (Neumann et al., 2012). In mouse cells, DSBs exhibiting increased mobility are the source of chromosomal translocations (Roukos et al., 2013). Thus, increased DNA mobility in response to DNA damage acts as a double-edged sword since it promotes homologous pairing but in some cases also it leads to potentially mutagenic DNA repair events.

As chromosome mobility is an important facet of the DNA damage response, investigating the nature of DNA diffusion in the context of repair is essential to understand how cells maintain their genome integrity. The mode of diffusion of a moving object drastically changes the way it explores the available space and the time to reach a specific target destination (Guerin et al., 2012). Thus, the kinetics of co-localization between two biological entities strongly depends on how these entities diffuse. One method to characterize DNA mobility consists of fluorescently marking chromosomal loci, measuring their displacement over time and calculating their mean square displacement (MSD) (Meister et al., 2010). The MSD curve represents the amount of space a locus has explored in the nucleus and its shape reveals the nature of DNA motion. When a particle freely diffuses, its MSD curve is linear with time and its motion is called “Brownian”. However, in living cells, DNA motion is often slower than Brownian diffusion and is called “sub-diffusive” (Barkai et al., 2012). Several types of sub-diffusive motion have been observed. When a chromosomal locus is confined inside a sub-volume of the nucleus, the motion is called *confined sub-diffusion* and the MSD exhibits a plateau (Marshall et al., 1997). When the force or structure that restricts the motion is not a simple confinement but is modulated in time and space with scaling properties, the motion is called *anomalous sub-diffusion* (Barkai et al., 2012; Metzler et al., 2014). In this case, sub-diffusive loci are constrained but unlike confined loci, they can diffuse without boundary and thus reach further targets if given enough time. For sub-diffusive motion, the MSD exhibits a power law (MSD ~ At^α^), where *α*, the anomalous exponent, is smaller than 1. The anomalous exponent *α* is linked to the degree of recurrence of DNA exploration, i.e., the number of times a DNA locus reiteratively scans neighboring regions before reaching a distant position (Ben-Avraham, 2000). When α is small, the locus explores recurrently the same environment for a long time, while a large α indicates that the locus is able to explore new environments often. The anomalous diffusion coefficient *A* represents the amplitude of DNA motion; it is proportional to the diffusion coefficient only in the case of normal diffusion (when α=1), which is rarely observed in biological systems (Barkai et al., 2012). Previous DNA mobility studies reported confined diffusion (Backlund et al., 2015; Cabal et al., 2006; Heun et al., 2001; Marshall et al., 1997; Masui et al., 2011; Taddei et al., 2006) while others have reported anomalous diffusion (Backlund et al., 2015; Burnecki et al., 2012; Cabal et al., 2006; Hajjoul et al., 2013; Lucas et al., 2014; Weber et al., 2010). These studies have been realized with different microscopy techniques and illumination settings. When studying the diffusion of a specific locus, the time scale at which the data is collected translates into the spatial scale of the exploration studied. So far, no consensus has yet been reached to describe the nature of DNA motion probably because different studies interrogate different spatiotemporal scales and therefore interrogate potentially different processes.

Here we have investigated DNA mobility at different time scales ranging from milliseconds to minutes using ultra-fast imaging in living *S. cerevisiae* cells. We observe that DNA motion is sub-diffusive at time scales ranging from milliseconds to a few minutes, with an anomalous exponent of 0.5 stable at multiple scales in diploid yeast. However, in response to DSB, DNA mobility is dramatically altered in a different manner depending on the time scales. Damaged DNA loci exhibit distinct anomalous regimes with increased mobility at large time scale but surprisingly reduced mobility at shorter time scales (less than 10 s). Importantly, the presence of distinct regimes of diffusions is not an intrinsic property of the damaged locus: in the presence of random DSBs, undamaged loci also exhibits increased mobility at large time scales and reduced mobility at short time scales. In light of polymers physics, such a pattern of dynamics has been predicted when chromatin persistence length globally increases, indicating that chromatin undergo a general stiffening in response to DSB. Rad51, a central protein of homologous recombination, is required for local and global changes in mobility/stiffening. We propose that global chromatin stiffening following DSBs facilitates nuclear exploration by the damaged DNA ends bound by the Rad51 protein.

## RESULTS

### DNA mobility exhibits anomalous regimes at different time scales

To characterize DNA dynamics at different time scales, we used diploid cells with a locus fluorescently marked by the insertion of a tet-Operator (*tetO*) array at *URA3* (chromosome V). This *tetO* array is bound by Tet-Repressors, which are fused to red fluorescent proteins (TetR-RFP). We also tagged a structural component of the spindle pole body (SPB) with yellow fluorescent protein, Spc110-YFP, to serve as a marker of the relative nuclear position and to correct for drifting during image acquisition for time intervals longer than 100 ms. Rad52, an essential homologous recombination protein, is fused to cyan fluorescent protein (CFP) to detect the presence of DSB (Fig. 1A and 1B) (Miné-Hattab and Rothstein, 2012). We selected early S-phase cells with a single SPB (approximately 10% of the total population), and where the *tetO*/TetR-RFP and Spc110-YFP foci are in the same focal plane. We recorded 2-dimensional movies of these cells (see supplementary movie 1). To avoid cells with spontaneous DNA damage, we ensured that they did not contain a Rad52 focus. Cells were imaged at three time scales:

i. 10 ms time intervals (5 ms exposure time in RFP, no YFP data were acquired since the global movement of the nucleus is negligible during this interval),
ii. 100 ms time intervals (50 ms exposure in RFP followed by the same exposure in YFP to track the *URA3* locus and the SPB respectively),
iii. 1000 ms time intervals (500 ms exposure in RFP followed by the same exposure time in YFP).

**Figure 1.**
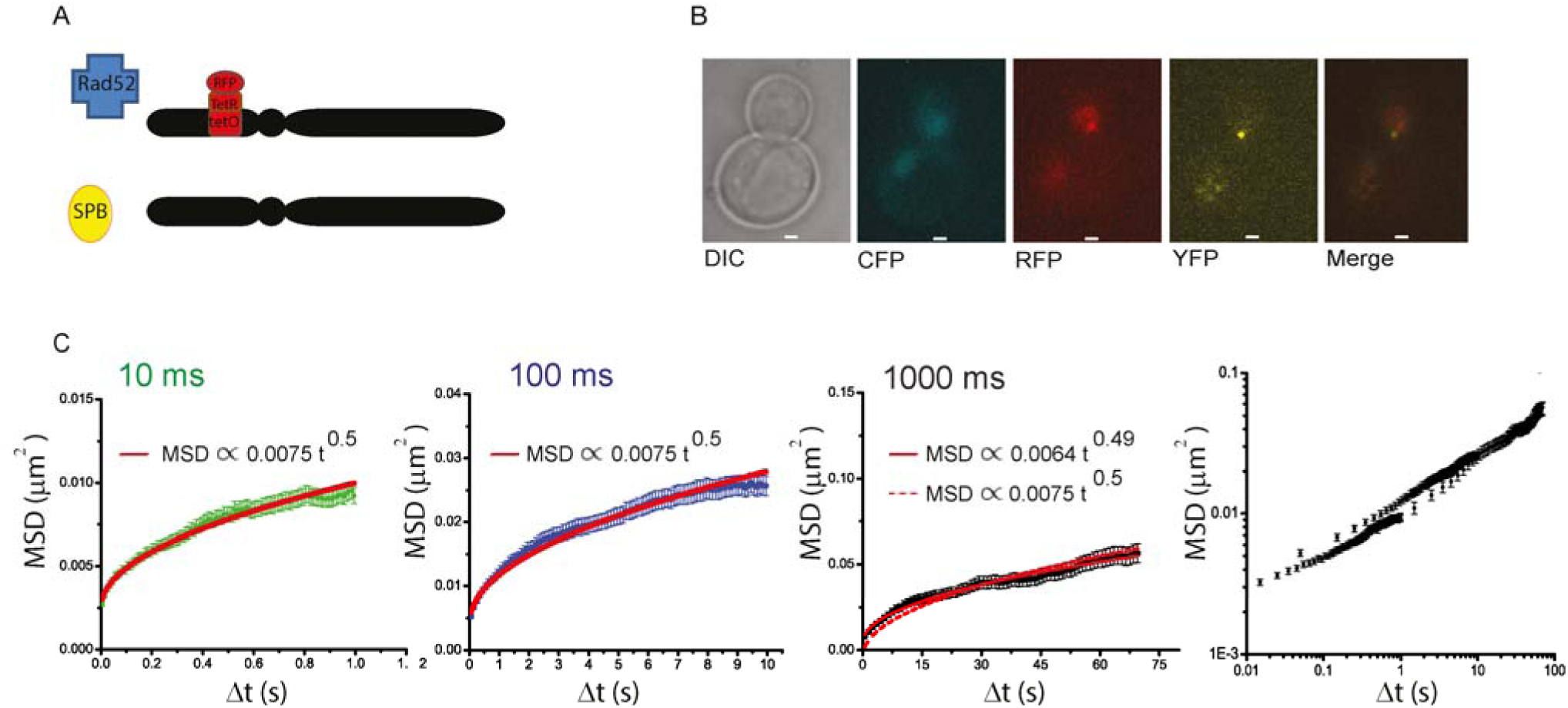
Mobility in diploid cells at different time scales. (A) Schematic of the strain: cells are diploids containing a *tet*O array (3x 112 copies) inserted at *URA3* (chromosome V). In addition, the Rad52, TetR and Spc110 proteins are tagged with CFP, RFP and YFP, respectively. Cells with the *tet*O locus and the SPB in the same focal plane are selected and the *tet*O locus is tracked in 2D over time using the SPB as a reference. (B) Typical transmitted, CFP, RFP, YFP and merge images of the cells used for the experiment. The experiment is performed on S-phase cells without a DSB as shown by the absence of a Rad52 focus. The scale bar is 1 μm. (C) Time and ensembled MSD of the *tet*O locus measured at 10 ms time intervals (first panel), 100 ms time intervals (second panel), 1000 ms time intervals (third panel). The fourth panel shows the merge of the 3 time scales plotted in log-log scale.

All acquisitions were performed with doses of light below the threshold of photo-toxicity (0.2 J/cm^2^, Logg et al., 2009). From the drift-corrected *x* and *y* coordinates of the *tetO*/TetR-RFP, we calculated the ensemble-averaged MSD using up to 1/3 of the shortest trajectory (which represents from 70 to 100 time points). Figure S1 shows that it is equivalent to measure the mobility at 100 ms time intervals using continuous illumination (50 ms exposure time in RFP followed by 50 ms in YFP) or discrete illumination (5 ms exposure time in RFP, followed by 5 ms in YFP and 90 ms of lag time). In addition, we mimicked acquisitions at 100 ms time intervals by averaging data acquired at 10 ms over 10 points and obtained similar MSD curves, allowing us to discard any photo-toxicity effects.

At the three time scales examined, the MSD curves show bending in linear scale suggesting sub-diffusion (Fig. 1C). The three MSDs curves are plotted together in log-log scale to allow their simultaneous visualization (Fig. 1C last panel). Since a power law curve can be mistaken for an exponential form that would account for confined diffusion, we used several methods to confirm the anomalous nature of DNA motion (Fig. S2). We then fitted our MSD curves with a power law. As experimental MSDs are altered by several artifacts (Backlund et al., 2015; Kepten et al., 2013), to fit MSD curves, we use an improved model that takes into account locus mobility during image acquisition and limited position accuracy, the latter (~ 90 nm) displaying little variation between conditions (see Material and Methods, Supplementary Text 1 and 2 and Fig. S3). Our approach is similar to the one described in Kepten *et al.,* but also includes exposure time as an additional parameter. In addition, we used the “multi-time scale fitting” approach described in Supplementary Text 2. For diploid cells, we obtain MSD(t) ~ 0.0075 t ^0.5^ at 10 and 100 ms time intervals (R2= 0.995 and 0.996). At large time scales (1000 ms time intervals), we observe a different anomalous diffusion coefficient *A* (MSD (t) ~ 0.0064 t ^0.49^, R2 = 0.992), likely due to a transition toward confined motion previously observed at larger time scales (10 s time intervals) (Miné-Hattab and Rothstein, 2012).

Since DNA mobility has been investigated in haploid cells using a multi time-scale approach (Hajjoul et al., 2013), we also measured DNA mobility in haploid cells containing a *tetO*/TetR-RFP array at *URA3* and the Spc110-YFP marked SPB (Fig. S4A and B). S-phase cells harboring a single SPB were imaged at the three time scales (10, 100 and 1000 ms time intervals). Similar to our observations in diploid cells, haploids follow sub-diffusive motion, however, unlike diploids, a 0.5 anomalous exponent is not seen at all time scales (Fig. S4C, D and E). Indeed, at 10 ms time intervals, we observe a different regime with MSD(t) ~ 0.011 t 0.38 (R2=0.999, Fig. S4C). Our results show a significant cell-to-cell variability (Fig. S5A, B and C) consistent with previous studies (Bronstein et al., 2009; Therizols et al., 2011). However, statistical analysis of the anomalous exponent (Fig. S5D, E and F) shows that the different exponents measured here are not due to insufficiencies in the dataset but rather reflect the existence of distinct anomalous regimes of the *URA3* locus depending on the time scale. Strikingly, a significant increase of the global MSD in haploids is seen compared to diploids at every exposure time investigated, as indicated by the higher values of the anomalous diffusion coefficient *A* in haploids (Fig. S4F).

Table 1 presents a summary of the results obtained in both diploids and haploids. Overall, the motion of the *URA3* locus is sub-diffusive for time scales ranging from milliseconds to minutes, and is well fit by a 0.5 anomalous exponent in most conditions. However, a universal regime is not sufficient to describe DNA motion, since at the very short time scale (10 ms), the anomalous exponent drops to 0.38 in haploids and at all-time scales examined, diploid cells exhibit lower MSD curves than haploids.

**TABLE 1:**
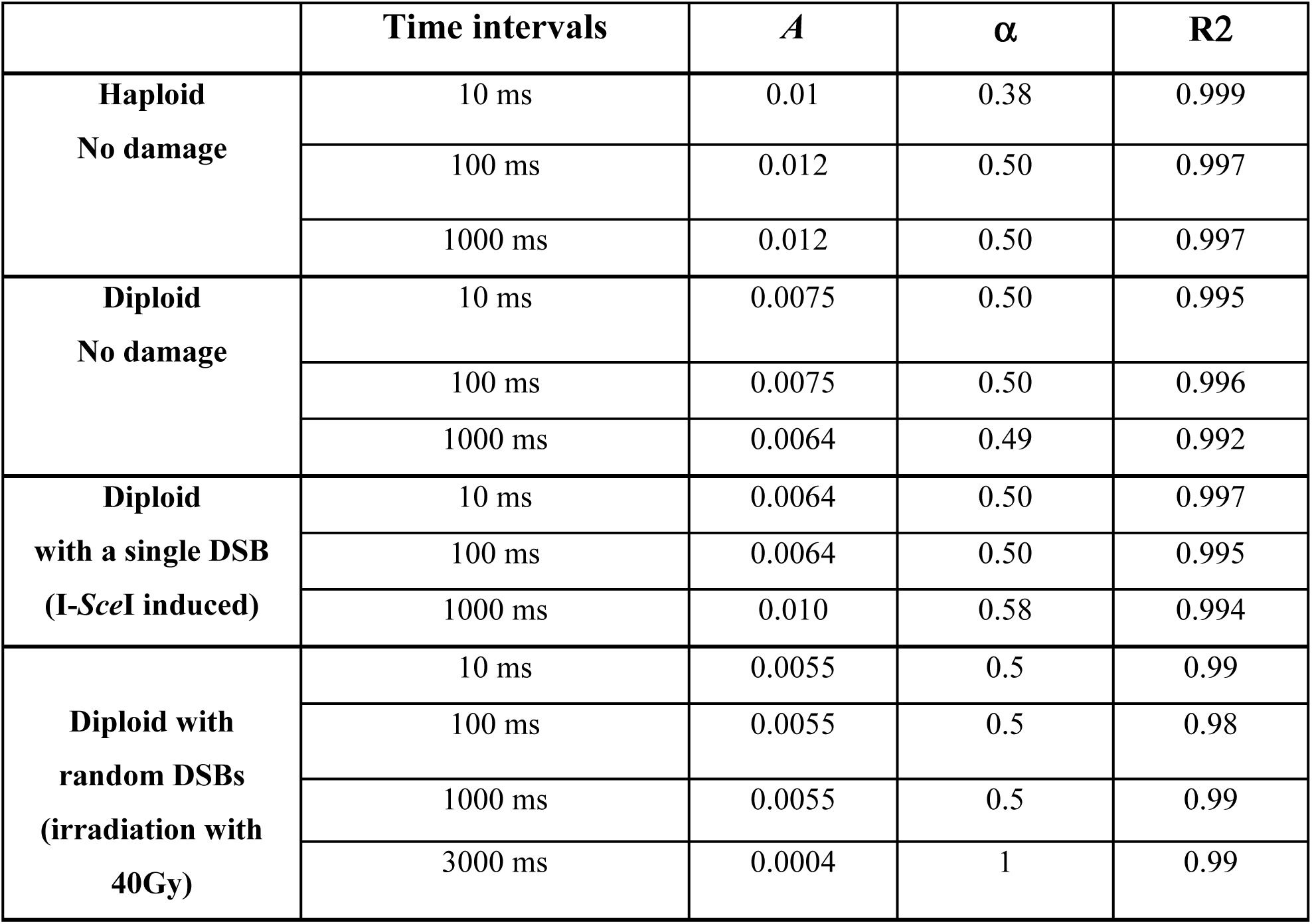
Summary of anomalous motion measurements. The anomalous diffusion coefficients *A* and the anomalous exponents α were obtained by fitting the MSD curves with *At^α^* + *b* (see equation 1 from Supplementary Text 1). The parameter *b* is an experimental offset allowing the correction of artifacts during the acquisition (i.e., localization accuracy and motion blur, see Supplementary Text 1). The fitting method is described in detail in Supplementary Text 2.

|  | **Time intervals** | **$A$** | **$\alpha$** | **R2** |
| --- | --- | --- | --- | --- |
| **Haploid**<br>**No damage** | 10 ms | 0.01 | 0.38 | 0.999 |
|  | 100 ms | 0.012 | 0.50 | 0.997 |
|  | 1000 ms | 0.012 | 0.50 | 0.997 |
| **Diploid**<br>**No damage** | 10 ms | 0.0075 | 0.50 | 0.995 |
|  | 100 ms | 0.0075 | 0.50 | 0.996 |
|  | 1000 ms | 0.0064 | 0.49 | 0.992 |
| **Diploid**<br>**with a single DSB**<br>**(I-SceI induced)** | 10 ms | 0.0064 | 0.50 | 0.997 |
|  | 100 ms | 0.0064 | 0.50 | 0.995 |
|  | 1000 ms | 0.010 | 0.58 | 0.994 |
| **Diploid with**<br>**random DSBs**<br>**(irradiation with**<br>**40Gy)** | 10 ms | 0.0055 | 0.5 | 0.99 |
|  | 100 ms | 0.0055 | 0.5 | 0.98 |
|  | 1000 ms | 0.0055 | 0.5 | 0.99 |
|  | 3000 ms | 0.0004 | 1 | 0.99 |

### At short time scales, damaged DNA is less mobile than in the absence of DSB

Previous studies have shown that the mobility of a damaged DNA locus increases when its motion is observed at 10 s time intervals in diploids (Miné-Hattab and Rothstein, 2012) and 1.5 s time intervals in haploids (Dion et al., 2012). However, mobility after DNA damage at shorter time scales has never been investigated. Here, we measure at multi time-scales the mobility of a single I-*Sce*I induced DSB in the same strains used in our previous study (Miné-Hattab and Rothstein, 2012) (Fig. 2). We used diploid cells containing the homologous *URA3* loci fluorescently marked with a *lacO*/LacI-YFP and a *tetO*/TetR-RFP array, respectively (Fig. 2A). To induce a single DSB, the strain contains an I-*Sce*I target site 4 kb from the *tetO*/TetR-RFP locus, as well as RAD52-CFP used as a marker for the presence of the DSB. Cells were incubated in 2% galactose for 90 minutes to induce the DSB and induction was stopped by adding 2% glucose. Note that the time of Rad52 focus formation cannot be precisely known; however, all cells were observed after the same incubation time (90 min), when Rad52 foci start to appear (Miné-Hattab and Rothstein, 2012). We next selected S-phase cells harboring a single SPB and a Rad52 focus colocalizing with the *URA3* locus (*tetO/*TetR-RFP) in the SPB focal plane (Fig. 2B). We verified that the two homologous *URA3* loci were unpaired by choosing cells with a distant *URA3* homologue (*lacO*/LacI-YFP). The *tetO* and SPB positions were measured over time in 2D at three time scales (10, 100 and 1000 ms) using the exact same illumination conditions than in the absence of DNA damage. We then calculated ensemble-averaged MSDs on these cells (Fig. 2C).

**Figure 2.**
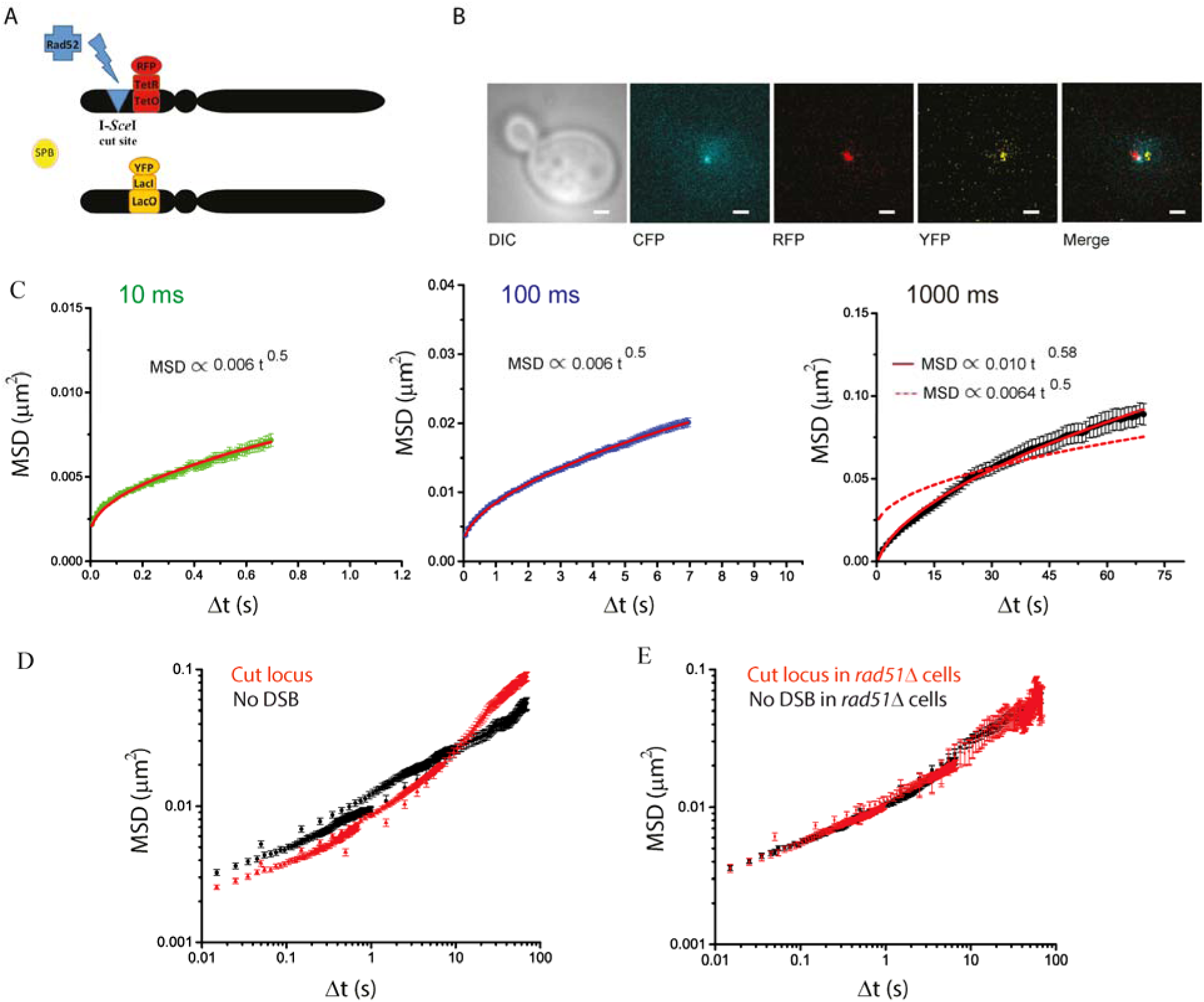
Mobility of a single I-*Sce*I induced DSB in diploid cells at different time scales. (A) Schematic of the strain used for the induction of <u>a single DSB</u> in diploid cells. Diploid cells contain both a *tet*O array (3x 112 copies) at *URA3* (chromosome V) and a *lac*O array (256 copies) at *URA3* on the other homologue. The Rad52, TetR and LacI proteins are fused with CFP, RFP and YFP, respectively. A single I-*Sce*I cut-site is located 4 kb from the *tetO* array. A galactose-inducible I-*Sce*I inserted at the *LYS2* locus allows regulated induction of a single DSB under galactose control (Miné-Hattab and Rothstein, 2012). (B) Typical transmitted, CFP, RFP, YFP and merge images of the cells after 90 minutes of galactose induction. Only S-phase cells containing a Rad52 focus co-localizing with the *tet*O locus and a distant *lac*O focus are selected. The scale bar is 1 μm. (C) Time and ensembled MSD of *tet*O array after induction of a single I-*Sce*I DSB. S-phase cells harboring a Rad52 focus colocalizing with the *tetO,* and a distant homologous locus (*lacO*-LacI-YFP) are selected. The mobility of the *tetO* array is measured at 10 ms time intervals. (left panel), 100 ms time intervals (middle panel) and 1000 ms (right panel). (D) Merge of the 3 time scales plotted in log-log scale, after I-*Sce*I induction (red curves). The MSDs of the *tetO* array in the absence of DNA damage is shown as a reference (black curve, same data as Fig. 1C, fourth panel). (E) MSDs of the *tetO* array after induction of a single I-*Sce*I DSB in *rad51Δ* cells. Mobility is measured at 10 ms, 100 ms and 1000 ms time intervals and the 3 MSD curves are shown in log-log scale. The mobility of the *tetO* array in the absence of damage in *rad51Δ* cells is also shown (black).

Similar to our results in the absence of DSBs, the damaged locus follows anomalous diffusion. At the longest time scale (1000 ms), we find the highest values of *A* and α in this study (MSD(t) ~ 0.010 t ^0.58^, R2 = 0.994) indicating increased mobility consistent with previous observations at 10 s time intervals (Miné-Hattab and Rothstein, 2012). Surprisingly, the motion of the damaged locus observed at shorter time intervals exhibits a lower amplitude and anomalous exponent than that seen in undamaged cells, signifying reduced mobility (MSD(t) ~ 0.0064 t ^0.50^, R2 = 0.995 and R2 = 0.997 at 100 and 10 ms time intervals respectively, Fig. 2C and Table 1). As a consequence, the MSD curve of the damaged locus crosses that of the undamaged one at time t~10s (Fig. 2D). Importantly, all cells are observed during the same stage of DNA repair, i.e., when the first Rad52 foci start to appear and homologous loci are unpaired. These cells are examined during 50 to 150 seconds depending on the time scale used: this acquisition time is negligible compared to the time necessary for DNA pairing in this system (Miné-Hattab and Rothstein, 2012). Thus, the differences in mobility observed here are likely solely due to the different time scales used to observe the damaged locus.

Overall, multi-time scales imaging of a damaged locus reveals that broken ends are less mobile at short time scales and exhibits different diffusion properties than at other scales. The reduced mobility seen at short time intervals reflects an important property of the damaged end, which might be triggered by the DNA repair machinery itself.

### Reduced mobility of the damaged locus at short time-scales is Rad51-dependent

Rad51, the central protein of HR, is required for increased mobility of a damaged end at large times scales (Dion et al., 2012; Miné-Hattab and Rothstein, 2012). To test whether *RAD51* is also involved in the reduced mobility that we observe at short time scales, we measured the mobility of a damaged end in *rad51*Δ diploid cells with the exact same illumination settings. Both alleles of the *RAD51* gene in the diploid strain described above were deleted. We then measured the *URA3* mobility in the absence of damage and after 90 minutes of DSB induction at 10, 100 and 1000 ms time intervals. Again, after DSB induction, we selected S phase-cells harboring a Rad52 focus colocalizing with the *URA3* locus, a single SPB and an unpaired distant *URA3* homologue. In the absence of *RAD51*, damaged loci failed to exhibit either increased or decreased mobility, indicating that Rad51 is required for changes in mobility at the damaged site at both short and long time scales (Fig. 2E).

### Random DSBs also provoke global reduced mobility at short time scales in a Rad51-dependent manner

An important question is whether changes in chromatin conformation and dynamics are localized around the site of damage or also affect the rest of the genome. In diploid yeast, increased mobility in response to DSB is not an intrinsic property of the damaged locus. Indeed, induction of DNA damage on a different chromosome or ionizing irradiation provokes global increased mobility affecting the whole genome (Miné-Hattab and Rothstein, 2012, 2013; Seeber et al., 2013). We thus tested whether reduced mobility over shorter time scales is restricted to the damaged end or can occur genome wide. We used diploid cells with a *tetO*/TetR-RFP at the *URA3* locus, Spc110-YFP and Rad52-CFP used as a marker for DSBs. We irradiated cells with 40 Gy, equivalent to 4 DSBs per nucleus in average (Ma et al., 2008). We then selected S-phase cells with a single SPB in the same focal plane as the *URA3* locus, as well as a Rad52 focus (Fig. 3A and B). Unlike the previous experiments, Rad52 foci do not colocalize with the *URA3* locus, and since chromosome V represents less than 2% of the genome, the probability of a DSB on that chromosome is extremely low. Immediately following irradiation, we measured the *URA3* mobility at four time scales (10, 100, 1000 and 3000 ms time intervals) and plotted the MSD curves (Fig. 3C and Fig. S6).

**Figure 3:**
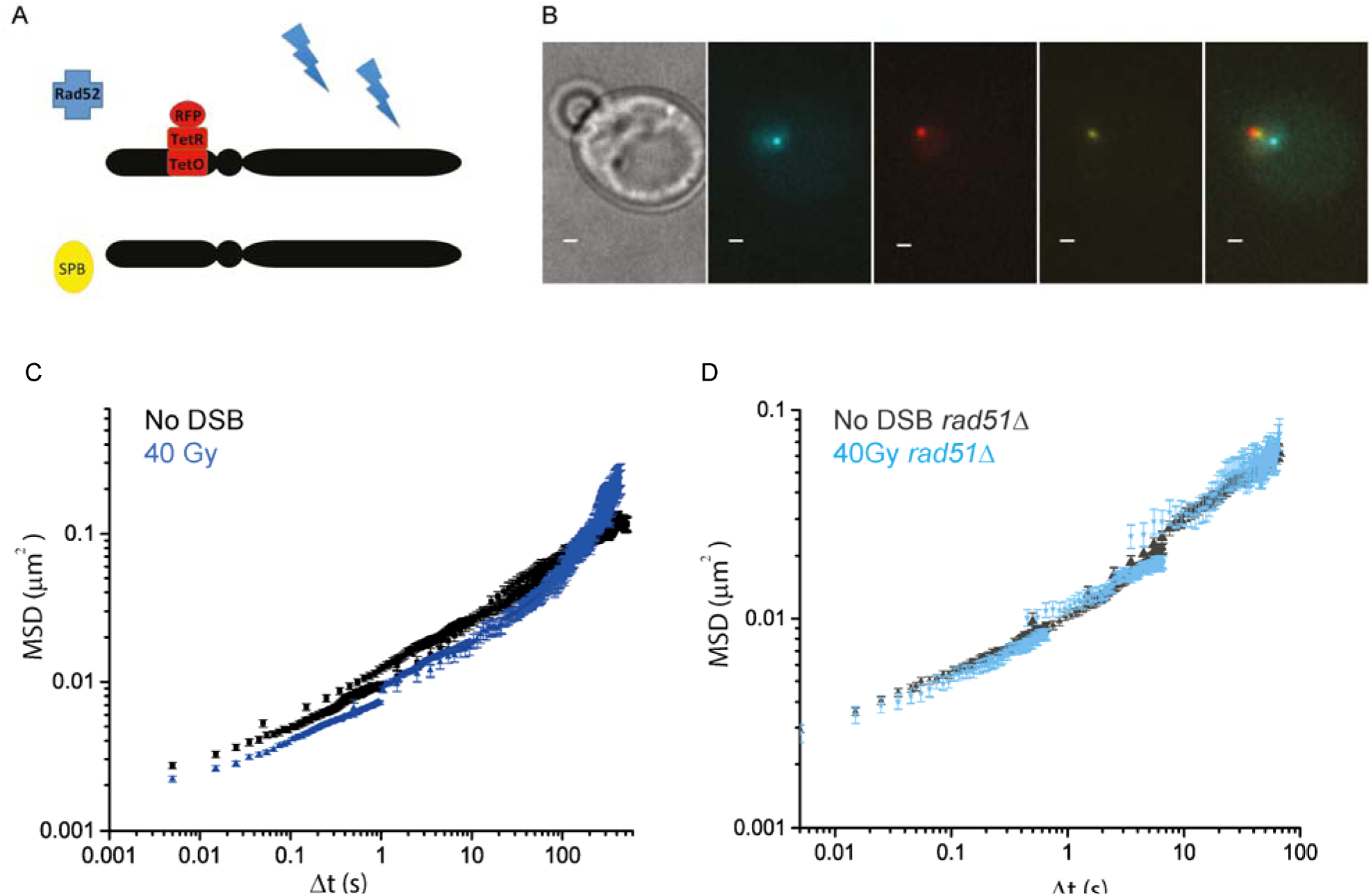
Mobility of the *URA3* locus in response to random DSBs (irradiation of 40 Gy). (A) Schematic of the strain: cells are diploids containing a *tet*O array (3x 112 copies) inserted at *URA3* (chromosome V). In addition, the Rad52, TetR and Spc110 proteins are tagged with CFP, RFP and YFP, respectively. Cells with the *tet*O locus and the SPB in the same focal plane are selected and the *tet*O locus is tracked in 2D over time using the SPB as a reference. (B) Typical transmitted, CFP, RFP, YFP and merge images of the cells after X-irradiation (40 Gy). Only S-phase cells containing a Rad52 focus are selected. Rad52 foci do not colocalize with the *tetO* array since DSBs are random. The scale bar is 1 μm. (C) MSD curves of the *tetO* array after 40 Gy measured at 10, 100 and 1000 ms time intervals (dark blue). The MSDs of the same locus in the absence of DNA damage is shown as a reference (black curve, same data as Fig. 1C, fourth panel). (D) MSDs of the *tetO* array after 40 Gy in *rad51Δ* cells. Mobility is measured at 10 ms, 100 ms and 1000 ms time intervals and the 3 MSD curves are shown in log-log scale (light blue). The mobility of the *tetO* array in the absence of damage in *rad51Δ* cells is also shown (black).

At the longest time scale (3000 ms), the MSD of irradiated cells is higher than in the absence of damage, consistent with global increased mobility previously reported at 10 s time intervals (Miné-Hattab and Rothstein, 2012). At 1000, 100 and 10 ms time intervals, undamaged loci exhibit a lower amplitude compared to that seen in the absence of DSB, signifying global reduced mobility at these three time scales (MSD(t) ~ 0.0055 t ^0.5^, R2 = 0.99, Fig. 3C and S6). As observed at the site of damage, both global increased and global reduced mobility are RAD51-dependent, since irradiated *rad51Δ* cells do not exhibit any increase in mobility (Fig. 3D).

## DISCUSSION

### Multi time-scale microscopy reveals the composite nature of DNA motion

Previous studies have shown that DNA motion observed at long time scales (1.5 to 10 s time intervals) is confined within a sub-nuclear volume (Heun et al., 2001; Marshall et al., 1997; Miné-Hattab and Rothstein, 2012). In response to DSBs, DNA becomes more mobile and explores a larger nuclear volume (Dion et al., 2012; Miné-Hattab and Rothstein, 2012). Here, by investigating DNA motion at time scales 10, 100 and 1000 times faster, we find that the MSD curves do not exhibit a plateau indicating that the effect of confinement is not observed at these scales. Instead, the MSD curves are in excellent agreement with a power law, MSD(t) ~ At^α^, a signature of anomalous diffusive motion (Barkai et al., 2012). At the longest time scale studied here (1000 ms time intervals), velocity auto-correlation functions indicate that the motion is not purely anomalous (Fig. S2D); this time scale likely corresponds to a transition toward confined motion, previously described at 10 s and larger time intervals (Marshall et al., 1997; Miné-Hattab and Rothstein, 2012). In addition, our data show a fundamental difference in DNA mobility between haploids and diploids (Fig. S4). The anomalous diffusion coefficient *A* is nearly two times higher in haploids indicating that the *URA3* locus is more mobile in haploids. One possible explanation for this difference is that the chromatin in haploid cells is less dense at the *URA3* locus.

In response to a single DSB, we find distinct anomalous regimes depending on the time scales, with both *A* and *α* increasing between the 100 ms and 1000 ms time intervals experiments (Fig. 2 and Table 1). These changes indicate a fundamental difference in the way the damaged locus explores the nuclear space at different time scales. The anomalous exponent α is linked to the degree of recurrence of the motion, low α corresponding to a locus that rescans neighboring loci many times in a highly recurrent manner (Ben-Avraham, 2000). The anomalous diffusion coefficient *A* reflects the amount of volume explored by a locus as a function of time (i.e., the amplitude of the motion). At 1000 ms time intervals, the higher *A* together with the higher *α* of the damaged locus indicates that the damaged locus moves with a larger amplitude and in a less redundant fashion (Table 1, A = 0.010, α = 0.58 after I*Sce*I induction compared to 0.0064 and 0.49 before damage). On the other hand, the low anomalous diffusion coefficient *A* of the damaged locus at 10 and 100 ms time intervals reflects reduced amplitude of the damaged DNA motion (Table 1, *A* = 0.0064 for the damaged DNA compared to 0.0075 in the absence of DSB). These changes in nuclear exploration at the broken end are Rad51-dependent, highlighting the role of Rad51 in regulating chromatin dynamics at the site of damage.

Importantly, the existence of different regimes of diffusion depending on the time scales is not an intrinsic property of the damaged end. In response to random DSBs (40 Gy), we also observe distinct anomalous regimes, with global increased mobility at long time scales and global reduced mobility at short time scales (Fig. 3C, Fig. S6 and Table 1). Interestingly, the MSD curves of cut loci *versus* loci in undamaged cells cross at 10 s (Fig. 2D) whereas the ones in irradiated cells *versus* undamaged cells cross at 100 s (Fig. 3C). In other words, it takes 10 s on average for a broken locus to cover larger distances than in the absence of a DSB, allowing the damaged site to reach further targets; in contrast, following random DSBs, the same locus needs 100 s on average to cover larger distances than in the absence of DSBs. Thus, upon DNA damage, changes in mobility have a stronger effect at the damaged locus than in the rest of the genome. This difference suggests that local and global changes in mobility are regulated differently. Overall, our findings show that a single mode of diffusion is not sufficient to describe DNA motion at different time scales. Instead, following DSBs, DNA motion is composed of several diffusion regimes that simultaneously drive DNA at each time scale. Such changes in the sub-diffusion mode dramatically modify the balance between surrounding and distant chromatin sampling. Thus, in the presence of DNA damage, the existence of multi time-scales regimes of diffusion may reflect changes in chromatin conformation that increase homology search efficiency.

### Origin of sub-diffusive motion

To link the anomalous diffusion measured here with possible changes in chromatin conformation, we discuss several models that explore the origin of anomalous diffusion to help gain insight into the biological consequences of multiple DNA diffusion regimes. Sub-diffusive motion has been observed in bacteria, yeast and human cells, with anomalous exponents ranging from 0.32 to 0.77 (Backlund et al., 2014; Bronstein et al., 2009; Hajjoul et al., 2013; Lucas et al., 2014; Weber et al., 2010). Although different time scales were examined in these studies, none clearly showed the existence of multiple anomalous regimes with the exception of a study by Bronstein *et al.* (Bronstein et al., 2009). In their study, they found that in U2OS cells, diffusion of telomeres is composed of two transient anomalous regimes and becomes close to Brownian at time scales greater than 5 minutes (Bronstein et al., 2009). Their work emphasizes the importance of the time scale of observation for interpreting DNA motion.

To understand the origin of anomalous sub-diffusion of DNA in the nucleus, several models have been proposed:

1. The nucleoplasm is modeled as a visco-elastic medium. Mathematical models of visco-elasticty are fractional Brownian motion (FBM) and fractional Langevin motion (FLM) (Metzler and Klafter, 2000). In these models, DNA is subjected to frictional forces that are not proportional to DNA velocity.
2. The nucleoplasm is modeled as a fractal. DNA loci are free explorers moving in a restricted geometry with scale-less properties imposed by nuclear crowding (Condamin et al., 2007).
3. The nucleoplasm is modeled as a polymer melt. DNA loci are represented by monomers whose motion is driven by the properties of this melt. Several polymer models have been suggested to describe DNA mobility, from diluted regimes (Rouse and Zimm models (Andrews, 2014; Vasquez and Bloom, 2014)) to a larger scale semi-diluted regime (“tube” model also called “blob” model (De Gennes, 1982)).

We did not consider the first two types of models because they do not predict distinct regimes. Since DNA is a polymer, we compared our experimental anomalous exponents to those obtained in the three polymer models referred to above. The “tube” model of De Gennes is the only one that predicts different regimes of anomalous diffusion arising in a polymer melt (De Gennes, 1982). This model predicts three anomalous diffusion regimes: the Rouse regime at a diluted scale (where *α* = 0.5), the “relaxation of the coil” regime (*α* = 0.25) followed by the “reptation” regime (*α* = 0.5) at a semi-diluted scale and concentration where adjacent chains constrain the motion. Finally, it predicts Brownian diffusion (*α* = 1) at the macroscopic scale. The reptation regime of this model is obtained by averaging different conformations of polymer entanglements, thus it is the only model that takes into account potential cell-to-cell variability (De Gennes, 1982). This regime has been directly observed *in vitro* for constrained DNA melts (Perkins et al., 1994); moreover, in live U2OS cell experiments, diffusion of telomeres observed at 1 s time intervals were explained by the reptation regime (Bronstein et al., 2009). Importantly, using the tube model, simulations of polymer dynamics show how global stiffening of polymers affect their dynamics: when the global stiffness of polymers increases, their MSD become lower at short time scales and higher at long time scales (Faller and Müller-Plathe, 2001). As a consequence, the MSD curves before and after global stiffening cross (Faller and Müller-Plathe, 2001).

Our data show that, in most conditions, the anomalous exponent is 0.5. To explain this behavior, we favor the “reptation regime” *versus* the Rouse regime for the following 4 reasons: i) we observe an anomalous diffusion coefficient *A* in haploids that is double that of diploids: in the reptation regime, this difference in *A* would be well justified by a lower level of chromatin entanglements in haploids compared to diploids, whereas the Rouse regime does not explain it, ii) at the shortest time interval, haploids exhibit a regime with an anomalous exponent lower than 0.5, which may be a transition from *the relaxation of the coil* regime to the reptation regime, iii) we observe high cell-to-cell variability, as predicted by the reptation regime, and more importantly, iv) modeling the global stiffening of polymers in the reptation regime leads to reduced mobility at short time scales and increased mobility at long scales (Faller and Müller-Plathe, 2001), which is consistent with our experimental observations after DNA damage (Fig. 2 and 3).

### Linking DNA mobility and chromatin persistence length

In relation with polymer physics, the changes in anomalous coefficient *A* that we measure reveal valuable information on chromatin plasticity in response to DSBs. Indeed, in the reptation regimes, where *α* = 0. 5, the anomalous diffusion coefficient *A* negatively correlates with the global chromatin persistence length (Faller and Müller-Plathe, 2001). To our knowledge, there is no consensus on the exact equation linking *A* and the persistence length *L_p_,* however, we can qualitatively discuss changes in global persistence length from our MSD measurements. In response to random DSBs, we measured the effect of DNA damage outside of the DSB, and found that DSBs provoke a genome wide effect at multi time scales. At small time scales, the anomalous diffusion coefficient *A* decreases resulting in crossing MSD curves between irradiated and undamaged cells (Fig. 3, Fig. S6 and Table 1). Such a decrease of *A* is predicted by the reptation regime when polymers become globally stiffer (Faller and Müller-Plathe, 2001). Thus, our results suggest that chromatin persistence length globally increases upon DSBs. Interestingly, global relaxation of chromatin in response to DSBs has been previously reported in mammalian cells (Ziv et al., 2006). In addition, recent studies in haploid yeast have shown that 20 to 40 % of the histones are degraded following DNA damage, and that the average distances between tagged loci increase after DNA damage (Hauer et al., 2017). Both of these studies can be interpreted as showing global chromatin decompaction upon DNA damage. Furthermore, the addition of nucleosomes on naked DNA to form a chromatin fiber is known to decrease its persistence length (Brower-Toland et al., 2002; Cui and Bustamante, 2000). Thus, the global increase in chromatin persistence length proposed here is consistent with the global chromatin relaxation and the degradation of histones reported after DNA damage. Finally, our results are in agreement with recent work from the Fabre laboratory in haploids yeast following zeocin-induced DSBs; global increased chromatin persistence length is thus a general chromatin response to DNA damage that is conserved between ploidies and types of damage.

As discussed above, changes in mobility have a stronger effect close to the damaged site compared to the rest of the genome (Fig. 2, 3 and S7). Unfortunately, we cannot easily interpret the MSD patterns observed close to the break in terms of persistence length, since the stiffening of a single chain in a polymer melt has not been investigated theoretically. However, *in vitro* measurements have already shown that the persistence length of the Rad51-ssDNA nucleo-filament itself is dramatically increased (Lee et al., 2013; Miné et al., 2007). Moreover, in the absence of the Rad51 protein, there is no change in mobility at both short and long time scales (Fig. 2E). Taken together, these findings underscore the role of Rad51 in local changes in stiffness at the DSB.

### The “needle in a decompacted ball of yarn” model

Using our observations, we formulate a model in which DSBs modify chromatin mobility both locally and globally, enhancing long distance explorations and minimizing local resampling. We propose that chromatin undergoes a genome-wide increase in persistence length in response to DSBs. Moreover, the damaged locus undergoes an additional effect due to the binding of repair proteins, making changes in mobility more pronounced at the broken locus. By stiffening the damaged DNA end, the repair machinery acts like a needle to help it search through the chromatin mesh, likened to a “ball of yarn” (Fig. 4). At short time scales, the stiffening of the Rad51-bound DNA leads to a reduction of mobility. However, concomitantly, the Rad51-DNA nucleo-filament enhances its ability to pass through the decompacted chromatin meshwork and escape adjacent obstacles more efficiently as observed at larger scales. Overall, global chromatin stiffening, combined with the “needle effect” at the DSB, facilitate the nuclear exploration of the damaged end throughout the genome, making it uniquely able to penetrate and explore the entangled chromatin network efficiently. Importantly, Rad51 is required for global changes in chromatin mobility/stiffness, indicating an essential role of the Rad51 proteins outside of the DSB in regulating global chromatin stiffening upon DNA damage. Given the role of checkpoint proteins and chromatin remodelers in haploid cells (Dion et al., 2012; Seeber et al., 2013), it will be interesting to elucidate their contribution to regulate chromatin flexibility in diploid yeast.

**Figure 4.**
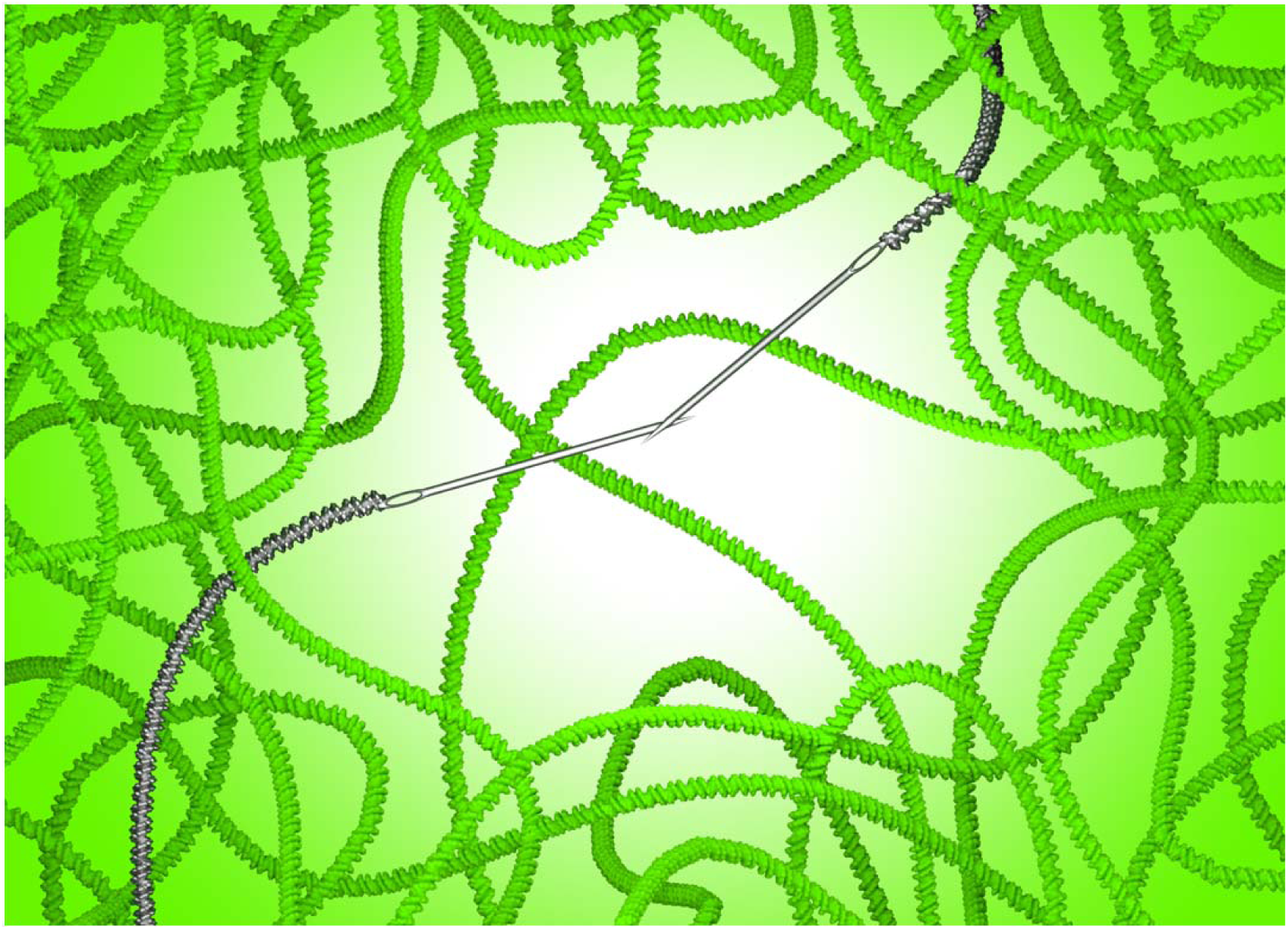
“Needle in a decompacted ball of yarn” model. To explain how chromatin exhibits decreased mobility at short time scales and simultaneously increased mobility at longer time scales after DNA damage, we propose the following model that we name “needle in a decompacted ball of yarn”. In response to double-strand breaks, chromatin undergoes a genome-wide increase in persistence length. In addition, the damaged locus undergoes an additional effect due to the binding of repair proteins, making changes in mobility more pronounced at the broken locus. Following a DSB, the repair complex forms a nucleofilament on the single strand DNA tail, which stiffens the damaged end (here shown as a needle), thus decreasing its mobility locally. However, the same stiffened DNA end acts like a “needle in a decompacted ball of yarn”, enabling it to escape adjacent obstacles more efficiently, thus increasing mobility at longer time scales. Chromatin is depicted as a green helical fiber. The two ends are represented as searching together consistent with earlier observations (Kaye et al., 2004; Lisby et al., 2003; Lobachev et al., 2004).

More generally, we propose that DNA diffusion is controlled by the local conformation of chromatin, i.e, the level of entanglement as described in the “tube model” (De Gennes, 1982). Future studies with super-resolution microscopy will allow the visualization of DNA organization at the nanoscale level *in vivo* to quantify how local and global DNA conformations are affected following damage (Recamier et al., 2014; Ricci et al., 2015). It will also be important to examine other biological processes at different time scales to see whether the multi-scale exploration of space that we observe here is a general property that allows a more efficient sampling of the environment.

## MATERIAL AND METHODS

### Strains

All strains used in this work are isogenic to *RAD5+* W303 derivatives (Zhao et al., 1998) (Supplementary Table S1).

### Cell culture and DSB induction

Before microscopy, cells were grown to early log phase in 4 ml cultures of SC medium + 100 mg l^−1^ adenine + 2% raffinose at 23°C overnight. In the morning, 2% galactose was added to the culture for 90 minutes to induce a single DSB at the I-*Sce*I cut-site. Cells were then pelleted, washed in SC + 100 mg l^−1^ adenine medium + 2% glucose to stop DSB induction and placed on a 1.4% agarose slab for microscopy (Miné-Hattab and Rothstein, 2012). During the DSB induction, the I-*Sce*I cutting starts to occur after 30 minutes of induction but the first Rad52 foci colocalizing with the *tetO* array are visible by microscopy after 90 minutes of induction and can last more than 30 minutes (Miné-Hattab and Rothstein, 2012). Thus, the time of DSB formation cannot be known precisely. To measure DNA mobility of the damaged locus, we select cells harboring a Rad52 focus after 90 minutes of induction. The DSB is not necessarily formed at the same time in all cells examined: however, since we measure time-ensemble averaged MSDs on several cells, the presence of distinct anomalous regimes does not reflect different mobilities dependent on time after DSB induction, but is solely due to the different time scales used to observe the locus.

### Irradiation

Cells analyzed by microscopy were pre-grown in SC + 100 mg/l of adenine + 2% glucose at 23°C overnight. Cells were then washed in the same media and exposed to 40 Gy of irradiation (X-rays,Yxlon International, Philipps). Irradiated cells were then immediately processed for imaging.

### Microscopy

Imaging of single DNA loci was performed on an inverted microscope Nikon Ti Eclipse (Nikon Instruments, Tokyo, Japan), with a high numerical aperture objective (1.49 NA) and 100X magnification. We used a perfect focus system (Nikon) designed to avoid drift on the Z-axis of the optical system and keep the cells in focus. The excitation laser beams (514 nm and 561 nm) were coupled to an optical fiber and focused in the back focal plane of the objective using an appropriate dual band dichroic (Semrock Di01-R488/561-25x36). Experiments were acquired with alternating pulsed excitation for the 514 nm laser and the 561 nm laser, respectively. Fluorescence emission from either YFP or RFP fluorescent proteins was filtered with a dual band emission filter with windows centered at 523 nm (45 nm bandpass) and 610 nm (45 nm bandpass) (Semrock FF01-523/610-25). The pixel size of the EMCCD is 16 μm (iXon 897, Andor Technology, Belfast, Ireland), producing a pixel size of 160 nm after magnification. For the experiments at 5 ms acquisition time, we imaged a small region of interest containing the cell of interest, which allowed us to obtain acquisition rates as fast as 200 Hz (5 ms per frame).

All the experiments have been acquired with doses of light inferior to 0.2 J/cm^2^, to protect cells from possible phototoxic effects (Logg et al., 2009). All the experiments have been performed in the same illumination conditions (see Results section and Fig. S1).

### Image analysis

We detected and tracked the *x* and *y* positions of the DNA loci using a home-made program inspired by the MTT tracking software (Serge et al., 2008). From the *x* and *y* positions of the DNA locus over time, we computed the time-ensemble averaged Mean Square Displacement (MSD) using the formula: MSD(τ) = <Δ**r**(τ)^2^> = <[**r**(*t*+τ) - **r**(*t*)]^2^>, where **r**(*t*) is the 2D-position of the particle at time *t*, and τ is the lag time between two positions of the particle used to calculate the displacement Δ**r**(τ) = **r**(*t*+τ) - **r**(*t*). The brackets, < >, are a time-average over *t* and over locus trajectories acquired from different cells. The error bar is the 95% confidence interval.

We confirmed the anomalous nature of the motion using several methods (Fig. S2). The theoretical MSD of sub-diffusive motion is a power law, MSD(t) = At^α^. However, experimental MSDs are altered from their theoretical shape due to various experimental limitations such as limited position accuracy or locus mobility during each image acquisition (Backlund et al., 2015; Kepten et al., 2013). Those limitations add an offset term, *b*, to the experimental asymptotic MSD, compared to the theoretical one: MSD*_experimental_*(t) = At^α^ + *b* (Hughes, 1995) (see Supplementary Text 1). Here, we derived a versatile formula of the MSD to compute this offset *b*, allowing us to accurately fit our experimental MSD derived from different acquisition schemes (see equation 1 from Supplementary Text 1, and Fig. S3). Our approach is more general than the one described in Kepten *et al*. (2013) since it also includes the exposure time as an additional parameter.

## AUTHOR CONTRIBUTIONS

JM-H: designed and performed the experiments, analyzed and interpreted the data, wrote the manuscript, coordinated the project.

VR: developed the improved fitting analysis of MSD curves, helped in the interpretation of the data.

II: worked on the conception of the microscope, helped in the interpretation the data, helped in writing the manuscript.

RR: helped in the interpretation of the data, co-wrote the manuscript.

XD: helped in the interpretation of the data.

## ACKOWLEDGMENTS

We would like to thank Leonid Mirny, Angela Taddei, Maxime Dahan and Yaron Shav-Tal for fruitful discussions about this work. We would like to thank also Eldad Kepten, Peter Thorpe and Chloé Guedj for fruitful comments on this manuscript. This work was funded by a Marie Curie International Outgoing Fellowship (JM-H), the ANR-12-PDOC-0035-01 (JM-H), the Fondation pour la Recherche Médicale (Foundation for Medical Research in France) (VR), National Institutes of Health (GM50237, GM67055 and GM118180 to RR).

## REFERENCES

Andrews, S.S. (2014). Methods for modeling cytoskeletal and DNA filaments. Physical biology 11, 011001.

Backlund, M.P., Joyner, R., and Moerner, W.E. (2015). Chromosomal locus tracking with proper accounting of static and dynamic errors. Physical review. E, Statistical, nonlinear, and soft matter physics 91, 062716.

Backlund, M.P., Joyner, R., Weis, K., and Moerner, W.E. (2014). Correlations of three-dimensional motion of chromosomal loci in yeast revealed by the double-helix point spread function microscope. Mol Biol Cell 25, 3619-3629.

Barkai, E., Garini, Y., and Metzler, R. (2012). Strange kinetics of single molecules in living cells. Physics today 65, 29-35.

Ben-Avraham, D.H. S. (2000). Diffusion and reactions in fractals and disordered systems Cambridge University Press.

Bronshtein Berger, I., Kepten, E., and Garini, Y. (2013). Single-Particle Tracking for Studying the Dynamic Properties of Genomic Regions in Live Cells. Methods Mol Biol 1042, 139-151.

Bronstein, I., Israel, Y., Kepten, E., Mai, S., Shav-Tal, Y., Barkai, E., and Garini, Y. (2009). Transient anomalous diffusion of telomeres in the nucleus of mammalian cells. Phys Rev Lett 103, 018102.

Brower-Toland, B.D., Smith, C.L., Yeh, R.C., Lis, J.T., Peterson, C.L., and Wang, M.D. (2002). Mechanical disruption of individual nucleosomes reveals a reversible multistage release of DNA. Proc Natl Acad Sci U S A 99, 1960-1965.

Burnecki, K., Kepten, E., Janczura, J., Bronshtein, I., Garini, Y., and Weron, A. (2012). Universal algorithm for identification of fractional Brownian motion. A case of telomere subdiffusion. Biophys J 103, 1839-1847.

Cabal, G.G., Genovesio, A., Rodriguez-Navarro, S., Zimmer, C., Gadal, O., Lesne, A., Buc, H., Feuerbach-Fournier, F., Olivo-Marin, J.C., Hurt, E.C., et al. (2006). SAGA interacting factors confine sub-diffusion of transcribed genes to the nuclear envelope. Nature 441, 770-773.

Chiolo, I., Minoda, A., Colmenares, S.U., Polyzos, A., Costes, S.V., and Karpen, G.H. (2011). Double-strand breaks in heterochromatin move outside of a dynamic HP1a domain to complete recombinational repair. Cell 144, 732-744.

Condamin, S., Benichou, O., and Klafter, J. (2007). First-passage time distributions for subdiffusion in confined geometry. Phys Rev Lett 98, 250602.

Cui, Y., and Bustamante, C. (2000). Pulling a single chromatin fiber reveals the forces that maintain its higher-order structure. Proc Natl Acad Sci U S A 97, 127-132.

De Gennes, P.G. (1982). Kinetics of diffusioncontrolled processes in dense polymer systems. II.Effects of entanglements. The Journal of Chemical Physics 76, 3322-3326.

Dimitrova, N., Chen, Y.C., Spector, D.L., and de Lange, T. (2008). 53BP1 promotes non-homologous end joining of telomeres by increasing chromatin mobility. Nature 456, 524-528.

Dion, V., Kalck, V., Horigome, C., Towbin, B.D., and Gasser, S.M. (2012). Increased mobility of double-strand breaks requires Mec1, Rad9 and the homologous recombination machinery. Nat Cell Biol 14, 502-509.

Faller, R., and Müller-Plathe, F. (2001). Chain stiffness intensifies the reptation characteristics of polymer dynamics in the melt. Chemphyschem: a European journal of chemical physics and physical chemistry 2, 180-184.

Guerin, T., Benichou, O., and Voituriez, R. (2012). Non-Markovian polymer reaction kinetics. Nature chemistry 4, 568-573.

Haber, J.E., and Leung, W.Y. (1996). Lack of chromosome territoriality in yeast: promiscuous rejoining of broken chromosome ends. Proc Natl Acad Sci U S A 93, 13949-13954.

Hajjoul, H., Mathon, J., Ranchon, H., Goiffon, I., Mozziconacci, J., Albert, B., Carrivain, P., Victor, J.M., Gadal, O., Bystricky, K., et al. (2013). High-throughput chromatin motion tracking in living yeast reveals the flexibility of the fiber throughout the genome. Genome research 23, 1829-1838.

Hauer, M.H., Seeber, A., Singh, V., Thierry, R., Sack, R., Amitai, A., Kryzhanovska, M., Eglinger, J., Holcman, D., Owen-Hughes, T., et al. (2017). Histone degradation in response to DNA damage enhances chromatin dynamics and recombination rates. Nat Struct Mol Biol 24, 99-107.

Heun, P., Laroche, T., Shimada, K., Furrer, P., and Gasser, S.M. (2001). Chromosome dynamics in the yeast interphase nucleus. Science 294, 2181-2186.

Hughes, B.D. (1995). Random Walks and Random Environments: Volume 1: Random Walks. Oxford science publications.

Jakob, B., Splinter, J., Conrad, S., Voss, K.O., Zink, D., Durante, M., Lobrich, M., and Taucher-Scholz, G. (2011). DNA double-strand breaks in heterochromatin elicit fast repair protein recruitment, histone H2AX phosphorylation and relocation to euchromatin. Nucleic Acids Res 39, 6489-6499.

Kaye, J.A., Melo, J.A., Cheung, S.K., Vaze, M.B., Haber, J.E., and Toczyski, D.P. (2004). DNA breaks promote genomic instability by impeding proper chromosome segregation. Curr Biol 14, 2096-2106.

Kepten, E., Bronshtein, I., and Garini, Y. (2013). Improved estimation of anomalous diffusion exponents in single-particle tracking experiments. Physical review. E, Statistical, nonlinear, and soft matter physics 87, 052713.

Lawrimore, J., Barry, T.M., Barry, R.M., York, A.C., Cook, D.M., Akialis, K., Tyler, J., Vasquez, P., Yeh, E., and Bloom, K. (2017). Microtubule dynamics drive enhanced chromatin motion and mobilize telomeres in response to DNA damage. Mol Biol Cell.

Lee, M., Lipfert, J., Sanchez, H., Wyman, C., and Dekker, N.H. (2013). Structural and torsional properties of the RAD51-dsDNA nucleoprotein filament. Nucleic Acids Res 41, 7023-7030.

Lisby, M., Antunez de Mayolo, A., Mortensen, U.H., and Rothstein, R. (2003). Cell cycle-regulated centers of DNA double-strand break repair. Cell Cycle 2, 479-483.

Lobachev, K., Vitriol, E., Stemple, J., Resnick, M.A., and Bloom, K. (2004). Chromosome fragmentation after induction of a double-strand break is an active process prevented by the RMX repair complex. Curr Biol 14, 2107-2112.

Logg, K., Bodvard, K., Blomberg, A., and Kall, M. (2009). Investigations on light-induced stress in fluorescence microscopy using nuclear localization of the transcription factor Msn2p as a reporter. FEMS yeast research 9, 875-884.

Lucas, J.S., Zhang, Y., Dudko, O.K., and Murre, C. (2014). 3D trajectories adopted by coding and regulatory DNA elements: first-passage times for genomic interactions. Cell 158, 339-352.

Ma, W., Resnick, M.A., and Gordenin, D.A. (2008). Apn1 and Apn2 endonucleases prevent accumulation of repair-associated DNA breaks in budding yeast as revealed by direct chromosomal analysis. Nucleic Acids Res 36, 1836-1846.

Marshall, W.F., Straight, A., Marko, J.F., Swedlow, J., Dernburg, A., Belmont, A., Murray, A.W., Agard, D.A., and Sedat, J.W. (1997). Interphase chromosomes undergo constrained diffusional motion in living cells. Curr Biol 7, 930-939.

Masui, O., Bonnet, I., Le Baccon, P., Brito, I., Pollex, T., Murphy, N., Hupe, P., Barillot, E., Belmont, A.S., and Heard, E. (2011). Live-cell chromosome dynamics and outcome of X chromosome pairing events during ES cell differentiation. Cell 145, 447-458.

Meister, P., Gehlen, L.R., Varela, E., Kalck, V., and Gasser, S.M. (2010). Visualizing yeast chromosomes and nuclear architecture. Methods Enzymol 470, 535-567.

Metzler, R., Jeon, J.H., Cherstvy, A.G., and Barkai, E. (2014). Anomalous diffusion models and their properties: non-stationarity, non-ergodicity, and ageing at the centenary of single particle tracking. Phys Chem Chem Phys 16, 24128-24164.

Metzler, R., and Klafter, J. (2000). The random walk’s guide to anomalous diffusion: a fractional dynamics approach. Physics Reports 339, 1-77.

Miné-Hattab, J., and Rothstein, R. (2012). Increased chromosome mobility facilitates homology search during recombination. Nat Cell Biol 14, 510-517.

Miné-Hattab, J., and Rothstein, R. (2013). DNA in motion during double-strand break repair. Trends Cell Biol 23, 529-536.

Miné, J., Disseau, L., Takahashi, M., Cappello, G., Dutreix, M., and Viovy, J.L. (2007). Real-time measurements of the nucleation, growth and dissociation of single Rad51-DNA nucleoprotein filaments. Nucleic Acids Res 35, 7171-7187.

Misteli, T. (2010). Higher-order genome organization in human disease. Cold Spring Harbor perspectives in biology 2, a000794.

Neumann, F.R., Dion, V., Gehlen, L.R., Tsai-Pflugfelder, M., Schmid, R., Taddei, A., and Gasser, S.M. (2012). Targeted INO80 enhances subnuclear chromatin movement and ectopic homologous recombination. Genes Dev 26, 369-383.

Perkins, T.T., Smith, D.E., and Chu, S. (1994). Direct observation of tube-like motion of a single polymer chain. Science 264, 819-822.

Recamier, V., Izeddin, I., Bosanac, L., Dahan, M., Proux, F., and Darzacq, X. (2014). Single cell correlation fractal dimension of chromatin: a framework to interpret 3D single molecule super-resolution. Nucleus 5, 75-84.

Ricci, M.A., Manzo, C., Garcia-Parajo, M.F., Lakadamyali, M., and Cosma, M.P. (2015). Chromatin fibers are formed by heterogeneous groups of nucleosomes in vivo. Cell 160, 1145-1158.

Roukos, V., Voss, T.C., Schmidt, C.K., Lee, S., Wangsa, D., and Misteli, T. (2013). Spatial dynamics of chromosome translocations in living cells. Science 341, 660-664.

Seeber, A., Dion, V., and Gasser, S.M. (2013). Checkpoint kinases and the INO80 nucleosome remodeling complex enhance global chromatin mobility in response to DNA damage. Genes Dev 27, 1999-2008.

Serge, A., Bertaux, N., Rigneault, H., and Marguet, D. (2008). Dynamic multiple-target tracing to probe spatiotemporal cartography of cell membranes. Nat Methods 5, 687-694.

Taddei, A., Van Houwe, G., Hediger, F., Kalck, V., Cubizolles, F., Schober, H., and Gasser, S.M. (2006). Nuclear pore association confers optimal expression levels for an inducible yeast gene. Nature 441, 774-778.

Therizols, P., Duong, T., Dujon, B., Zimmer, C., and Fabre, E. (2011). Chromosome arm length and nuclear constraints determine the dynamic relationship of yeast subtelomeres. PNAS 107, 2025–2030.

Vasquez, P.A., and Bloom, K. (2014). Polymer models of interphase chromosomes. Nucleus 5, 376-390.

Weber, S.C., Spakowitz, A.J., and Theriot, J.A. (2010). Bacterial chromosomal loci move subdiffusively through a viscoelastic cytoplasm. Phys Rev Lett 104, 238102.

Zhao, X., Muller, E.G., and Rothstein, R. (1998). A suppressor of two essential checkpoint genes identifies a novel protein that negatively affects dNTP pools. Mol Cell 2, 329-340.

Ziv, Y., Bielopolski, D., Galanty, Y., Lukas, C., Taya, Y., Schultz, D.C., Lukas, J., Bekker-Jensen, S., Bartek, J., and Shiloh, Y. (2006). Chromatin relaxation in response to DNA double-strand breaks is modulated by a novel ATM- and KAP-1 dependent pathway. Nat Cell Biol 8, 870-876.

